# Sexual conflict in self-fertile hermaphrodites: reproductive differences among species, and between individuals versus cohorts, in the leech genus *Helobdella* (Lophotrochozoa; Annelida; Clitellata; Hirudinida; Glossiphoniidae)

**DOI:** 10.1101/401901

**Authors:** Roshni G. Iyer, D. Valle Rogers, Christopher J. Winchell, David A. Weisblat

**Affiliations:** Dept. of Molecular & Cell Biology, 385 LSA, Univ. of California, Berkeley, CA 94720-3200, USA; Dept. of Electrical Engineering & Computer Sciences, Univ. of California, Berkeley, CA 94720-3200, USA

## Abstract

Leeches and oligochaetes comprise a monophyletic group of annelids, the Clitellata, whose reproduction is characterized by simultaneous hermaphroditism. While most clitellate species reproduce by cross-fertilization, self-fertilization has been described within the speciose genus *Helobdella*. Here we document the reproductive life histories and reproductive capacities for three other *Helobdella* species. Under laboratory conditions, both *H. robusta* and *H. octatestisaca* exhibit uniparental reproduction, apparently reflecting self-fertility, and suggesting that this trait is ancestral for the genus. However, the third species, *H. austinensis*, seems incapable of reproduction by self-fertilization, so we inferred its reproductive life history by analyzing reproduction in breeding cohorts. Comparing the reproductive parameters for *H. robusta* reproducing in isolation and in cohorts revealed that reproduction in cohorts is dramatically delayed with respect to that of isolated individuals, and that cohorts of leeches coordinate their cocoon deposition in a manner that is not predicted from the reproductive parameters of individuals reproducing in isolation. Finally, our comparisons of reproductive capacity for individuals versus cohorts for *H. robusta*, and between different sizes of cohorts for *H. austinensis*, reveal differences in resource allocation between male and female reproductive roles that are consistent with evolutionary theory.

## INTRODUCTION

Leeches comprise a monophyletic group of segmented worms within the phylum Annelida. They occupy primarily freshwater habitats, as fluid-feeding ecto-parasites on vertebrate hosts, or predators or scavengers of freshwater invertebrates (Sawyer 1986). Molecular evidence indicates that leeches evolved from within the oligochaete annelids; together, these two taxa comprise the monophyletic assemblage of clitellate annelids (Kuo 2017; Ocegueara-Figueroa et al. 2016; Struck et al. 2011; Weigert et al. 2014; Zrzavy et al. 2009). Compared to oligochaetes, leeches are characterized by having lost segmentally iterated bristles (chaetae), by having a fixed number of segments, and by the presence of anterior and posterior suckers used for feeding and locomotion.

Certain leech species, primarily of the genus *Hirudo*, have proved valuable for analyzing neural circuits and behavior in terms of the activity and connectivity of individually identified neurons and for studies of individually defined neural cell types in culture (Muller et al. 1981). Other species, primarily in the family Glossiphoniidae, have been used in studies of cell lineage and embryonic development, speciation, predator-prey interactions and genome evolution in the super-phylum Lophotrochozoa (Blinn and Davies, 1989; Davies and Kasserra 1989; Weisblat and Kuo 2014; Sawyer 1986; Simakov et al. 2013). Thus, leeches generally, and those species in the glossiphoniid genus *Helobdella* in particular, provide models for integrating the questions and approaches from a wide range of biological sub-disciplines, from physiology and development to ecology, genomics and evolution in a less well explored branch of animals.

Leeches of the genus *Helobdella* are medium-sized (typically 1-3 cm as adults), neutrally pigmented, unobtrusive clitellate annelids, preying or scavenging on other invertebrates in shallow freshwater habitats. Molecular-phylogenetic analyses have revealed a surprising diversity of this genus: more than 50 species to date, many of which are difficult to distinguish morphologically (Bely and Weisblat 2006; Oceguera-Figueroa et al. 2010).

In the course of ongoing studies using different *Helobdella* species for studying embryonic development in Lophotrochozoa/Spiralia, we have observed differences in reproduction, feeding, and other behaviors. We have previously described the reproductive life history of a self-fertile *Helobdella* species identified as *H. triserialis* (Wedeen et al. 1990). The data presented here details our findings concerning the reproductive life history, under similar conditions of laboratory culture, for *H. robusta* (Shankland et al. 1992) and for a scute-bearing (*H. stagnalis*-like) species that we identify as *H. octatestisaca* (Lai et al. 2009) on the basis of its cytochrome oxidase 1 (CO1) sequence. Like *H. triserialis*, both of these species are self-fertile, as has also been reported for other glossiphoniid and piscolid species (Kua et al. 2010; Tan et al. 2004; Whitman 1878). Thus, we were surprised to discover that a third species, *H. austinensis* (Kutschera et al. 2013), is incapable of reproduction by self-fertilization. For this species we therefore inferred the reproductive life history of individuals by analyzing reproduction in breeding cohorts. This led us to compare the reproductive parameters for *H. robusta* raised in isolation and in cohorts, which yielded another surprising result. We found that reproduction by cohorts of *H. robusta* is dramatically delayed with respect to that of isolated individuals, and that cohorts of leeches coordinate their cocoon deposition in a manner that is not predicted from the reproductive parameters of isolated individuals.

## MATERIALS AND METHODS

### Animals

The taxonomy of the genus *Helobdella* is in flux, due in large part to the increased resolution provided by the advent of molecular sequence comparisons. Data presented here represent three operational taxonomic units (OTUs; Figure 01):

OTU1 is the recently described *H. austinensis* (Hau; Kutschera et al. 2013), collected from the wild in Austin, TX, and in continuous laboratory culture since the 1980s. OTU2 is *H. robusta* (Hro; Shankland et al. 1992), re-collected from its type location in Sacramento, CA. OTU3, collected from the same location as OTU2, is a *H. stagnalis*-like species, as defined by the presence of a nuchal scute on the dorsal surface at the boundary between the rostral and midbody segments. Molecular phylogenies have revealed that the morphologically defined *H. stagnalis* is in fact a complex of species (Beresic-Perrins 2017; Oceguera-Figueroa et al. 2010; Saglam et al., 2018); CO1 sequencing indicates that the species used here is *H. octatestisaca* (Hoc; Lai et al. 2014).

For molecular identification, the cytochrome oxidase 1 (CO1) sequences for the species used here are available as GenBank accession numbers: Hau, MH729328; Hro, MH729330; Hoc, MH729329). For comparison, we also discuss previously published data on reproductive life history from a fourth OTU, *H. triserialis* (Htr), originally collected in San Francisco, CA in the 1970s; this species was maintained in laboratory culture for several years through the early 1980s, but was lost from the laboratory and disappeared from its original location prior to the advent of molecular barcoding.

### Reproductive analysis

For some experiments, individuals for which the exact birthdate (defined here as the date of zygote deposition) was known were reared in isolation from early stages of development in small petri dishes (35 or 50 mm diameter), with daily feeding and changes of water (1/100 dilution of artificial seawater; Salinity for Reefs, Aquavitro) at room temperature (21-23°C). For other experiments, groups of late stage embryos or early juveniles, from a clutch for which the exact birth date was known, were isolated and reared as freely breeding cohorts, maintained as above except for being transferred as adults to larger containers (0.5-1 liter capacity pyrex bowls). With rare exceptions, animals in both conditions were checked daily for reproductive activity and deaths; embryos were usually removed and counted within 24 hours of zygote deposition. On occasions where clutch deposition was not observed immediately, the date of laying was estimated from the stage of development attained when the clutch was removed.

## RESULTS

### Reproductive life histories of individuals raised in isolation reveal differences among three self-fertile *Helobdella* species

Sexual reproduction by simultaneous hermaphrodites is the presumed ancestral state of clitellate annelids, although some species now rely in part or entirely on various modes of asexual reproduction (Zattara and Bely 2011, 2013); the ecology of sexual reproduction has been reviewed (Lively and Morran 2014). Cross-fertilization is required for most clitellate annelids, but several species have the ability to produce viable embryos without ever having had contact with prospective mates. While the possibility of other uniparental modes of reproduction has not been rigorously excluded, we observe polar body formation, indicative of maternal meiosis, in uniparental zygotes, and this capacity for reproducing without mating in leeches has generally been accepted as self-fertilization. Thus, we will use that term here. By this criterion, self-fertilization among leeches has been documented previously for the piscicolid species *Zeylanicobdella arugamensis* (Kua et al. 2010), and for at least three glossiphoniid species, including the species referred to by Whitman as *Clepsine marginata* (Whitman 1878), *Helobdella triserialis* (Wedeen et al. 1990) and *H. papillornata* (Tan 2004). Here, we document that the strains of *H. robusta* and *H. octatestisaca* that we have studied are also self-fertile, as well as being capable of cross-fertilization. Individuals raised in isolation from embryonic stages routinely produce viable young, and these progeny are also self-fertile when reared in isolation. In these self-fertilizing animals, we found no evidence of the externally implanted spermatophores that are seen upon cross-fertilization in these species. Thus, we conclude that self-fertilization does not involve implantation of a spermatophore, but rather is achieved internally.

For these self-fertile species, as for *H. triserialis* (Wedeen et al 1990), it was possible to directly measure the reproductive capacity (defined here as the number of young produced during the life of one individual) under defined conditions by rearing individuals in isolation from early stages of development until their death, removing and determining the size of all clutches for each individual; for comparison, previously published comparable data for *H. triserialis* (Wedeen et al 1990) is summarized here as well.

Previous work had shown that, when reared in isolation under laboratory conditions at room temperature, *H. triserialis* exhibits an egg-to-egg generation of time of about 70 days, then generates five clutches of embryos at 30-35 day intervals. Of these five clutches, the first and last were smaller and the third was the largest (Supplemental Table 1). For *H. triserialis* reared in isolation under these conditions, five clutches was a hard maximum; at least one individual survived for over three months after depositing its fifth clutch, well beyond the average inter-clutch interval, without further reproduction, and the production of fewer than five clutches was invariably associated with premature death of the animal. The average reproductive capacity measured for *H. triserialis* in those experiments was 302 offspring per individual.

**Table 1.** Clutch size data.

| Clutch number (S)* | Clutch size | Clutch size (min, max) |
| --- | --- | --- |
| C1 (10) | 20.3 +/- 4.1 | (15, 25) |
| C2 (14) | 51.1 +/- 16.3 | (3, 70) |
| C3 (14) | 71.1 +/- 19.5 | (34, 99) |
| C4 (11) | 77.6 +/- 19.9 | (55, 104) |
| C5 (11) | 69.1 +/- 23.5 | (30, 105) |
| C6 (9) | 51.0 +/- 30.7 | (7, 95) |
| C7 (6) | 44.2 +/- 27.3 | (12, 81) |
| C8 (1) | 13.0 | 13 |

***H. octitestisaca*: self-fertilizing, bloodworm diet (N = 5)**
| Clutch number (S)* | Clutch size | Clutch size (min, max) |
| --- | --- | --- |
| C1 (5) | 26.4 +/- 7.5 | (17, 37) |
| C2 (5) | 50.2 +/- 12.7 | (33, 66) |
| C3 (3) | 70.3 +/- 4.7 | (65, 74) |

***H. austinensis*: interbreeding cohort, bloodworm diet (N = 23)\*\***
| Clutch number (S)* | Clutch size | Clutch size (min, max) |
| --- | --- | --- |
| C1 (23) | 94.2 +/- 41.0 | (45, 179) |
| C2 (16) | 87.4 +/- 36.2 | (22, 160) |
| C3 (1) | 26.0 | (26, 26) |

***H. austinensis*: interbreeding cohort, bloodworm diet (N = 60)\*\***
| Clutch number (S)* | Clutch size | Clutch size (min, max) |
| --- | --- | --- |
| C1 (60) | 43.8 +/- 20.8 | (6, 117) |
| C2 (54) | 41.5 +/- 22.4 | (7, 93) |
| C3 (31) | 38.9 +/- 25.2 | (6, 97) |

***H. robusta*: interbreeding cohort, snail diet (N = 48)\*\***
| Clutch number (S)* | Clutch size | Clutch size (min, max) |
| --- | --- | --- |
| C1 (48) | 24.0 +/- 10.0 | (11, 65) |
| C2 (34) | 44.6 +/- 16.8 | (15, 85) |
| C3 (28) | 68.4 +/- 18.4 | (21, 102) |
| C4 (26) | 75.3 +/- 18.5 | (29, 114) |
| C5 (13) | 48.1 +/- 22.3 | (17, 85) |

Clutch size is average +/-standard deviation; \* Sample size (S); \*\*For interbreeding cohorts, clutch number was inferred as described in text.
***H. robusta*: self-fertilizing, snail diet (N = 16)**

The reproductive life history for *H. robusta* (summarized in Tables 1 and 2; for more detailed information see Supplemental Table 2) differs both qualitatively and quantitatively from that of *H. triserialis* under similar conditions. Firstly, the average egg-to-egg generation time for this species was 57 days and the average inter-clutch interval was less than 30 days. In addition, this species was capable of laying more than five clutches of embryos; a maximum of eight was observed. This increase in the number of clutches was associated with somewhat smaller clutch sizes, and with a more uniform distribution of clutch sizes (compare Table 2, Supplemental Tables 1 and 2). The average reproductive capacity for *H. robusta* raised in isolation was 267 offspring per individual, with a maximum of 392.

**Table 2.** Clutch interval data.

| Interval (S)* | Zygote-clutch interval | Inter-clutch interval | Inter-clutch interval (min, max) |
| --- | --- | --- | --- |
| **ZD-C1 (15)** | 56.3 +/- 8.7 |  | (37, 68) |
| **C1-C2 (15)** | 83.5 +/- 7.7 | 28 | (72, 93) |
| **C2-C3 (14)** | 107.4 +/- 10.5 | 23 | (92, 121) |
| **C3-C4 (12)** | 136.8 +/- 9.9 | 30 | (117, 150) |
| **C4-C5 (10)** | 170.4 +/- 16.0 | 33 | (142, 193) |
| **C5-C6 (8)** | 199.4 +/- 17.2 | 29 | (169, 222) |
| **C6-C7 (5)** | 243.3 +/- 29.6 | 44 | (217, 295) |
| **C7-C8 (1)** | 264.0 | 21 | 264 |

***H. octitesticata*: self-fertilizing, bloodworm diet (N = 5)**
| Interval (S)* | Zygote-clutch interval | Inter-clutch interval | Inter-clutch interval (min, max) |
| --- | --- | --- | --- |
| **ZD-C1 (5)** | 140.0 +/- 25.8 |  | (120, 180) |
| **C1-C2 (4)** | 160.8 +/- 37.7 | 21 | (137, 227) |
| **C2-C3 (3)** | 220.8 +/- 43.2 | 60 | (191, 270) |

***H. austinensis*: interbreeding cohort, bloodworm diet (N = 23)\*\***
| Interval (S)* | Zygote-clutch interval | Inter-clutch interval | Inter-clutch interval (min, max) |
| --- | --- | --- | --- |
| **ZD-C1 (23)** | 109.3 +/- 20.4 |  | (80, 148) |
| **C1-C2 (16)** | 188.4 +/- 36.6 | 79 | (148, 240) |
| **C2-C3 (1)** | 243.0 | 55 | 243 |

***H. austinensis*: interbreeding cohort, bloodworm diet (N = 60)\*\***
| Interval (S)* | Zygote-clutch interval | Inter-clutch interval | Inter-clutch interval (min, max) |
| --- | --- | --- | --- |
| **ZD-C1 (60)** | 109.7 +/- 19.3 |  | (70, 143) |
| **C1-C2 (54)** | 189.8 +/- 28.3 | 80 | (144, 234) |
| **C2-C3 (31)** | 273.1 +/- 57.2 | 83 | (237, 346) |

| Interval (S)* | Zygote-clutch interval | Inter-clutch interval | Inter-clutch interval (min, max) |
| --- | --- | --- | --- |
| **ZD-C1 (48)** | 107.4 +/- 10.7 |  | (94, 134) |
| **C1-C2 (34)** | 146.8 +/- 11.1 | 40 | (135, 170) |
| **C2-C3 (28)** | 184.9 +/- 11.0 | 48 | (171, 204) |
| **C3-C4 (26)** | 221.1 +/- 13.2 | 36 | (204, 241) |
| **C4-C5 (13)** | 252.8 +/- 9.2 | 32 | (241, 276) |

\* Sample size (S); Intervals given as average +/- standard deviation; \*\*For interbreeding cohorts, clutch intervals were inferred as described in text.

The reproductive life history we observed for isolated, self-fertilizing *H. octatestisaca*(summarized in Tables 1 and 2; more detailed information in Supplemental Table 3) differed markedly from those described for either *H. robusta* or *H. triserialis*. From among a sample of five individuals, the egg-to-egg generation time was longer (140 days versus 56 days for *H. robusta*), no individual produced more than three clutches of embryos, and the average reproductive capacity was dramatically less (119 versus 267 for *H. robusta*). Although we cannot exclude the possibility that these differences reflect culture conditions that were sub-optimal for *H. octatestisaca*, the average life-span of *H. octatestisaca* in this experiment (246 days) was not less than that of *H. robusta* under similar conditions (229 days), and several of the animals survived after laying their last clutch of embryos for periods of time that were much longer than the average inter-clutch interval. The difference in the degree of iteroparity between *H. robusta* and *H. octatestisaca* is also consistent with the fact that, within the genus *Helobdella*, molecular phylogenies place *H. robusta* and *H. triserialis* in one species complex, the “triserialis” series, and most of the scute-bearing species, including *H. octatestisaca* into a paraphyletic “stagnalis” series (Oceguera-Figueroa et al., 2010; Saglam et al. 2018).

### *Helobdella austinensis* does not reproduce in isolation

Given that *H. triserialis, H. papillornata, H. robusta* and *H. octatestisaca* are all self-fertile, it came as a surprise that we were unable to observe self-fertilization for *H. austinensis*, which is more closely related to *H. robusta* than are the other three self-fertile species of *Helobdella*. Individuals reared in isolation for months failed to reproduce, but soon became gravid when placed with other individuals (Michelle Levine, personal communication).

Thus, to study the reproductive life history of *H. austinensis*, we raised cohorts of interbreeding animals, tracking survivorship and reproductive activity of the adult cohort by noting the dates of deaths and clutch depositions, and the size of clutches produced during the collective life of the cohort, respectively.

### Life history analyses for cohorts of interbreeding *H. austinensis* suggest differences in reproductive behaviors from other *Helobdella* species

In two experiments, we studied cohorts of *H. austinensis* reared on a diet of commercially available “bloodworms” (frozen midge larvae); clutches of embryos were removed and counted as soon as they were observed, usually within 24 hours of having been deposited. The two cohorts, starting with 23 and 60 individuals, respectively, produced a total of 40 and 145 clutches, respectively. For each experiment, we tracked the number of surviving leeches within the cohort, the number of clutches deposited, the number of embryos per clutch and the aggregate number of embryos produced (Figures 2, 3).

**Figure 1.**
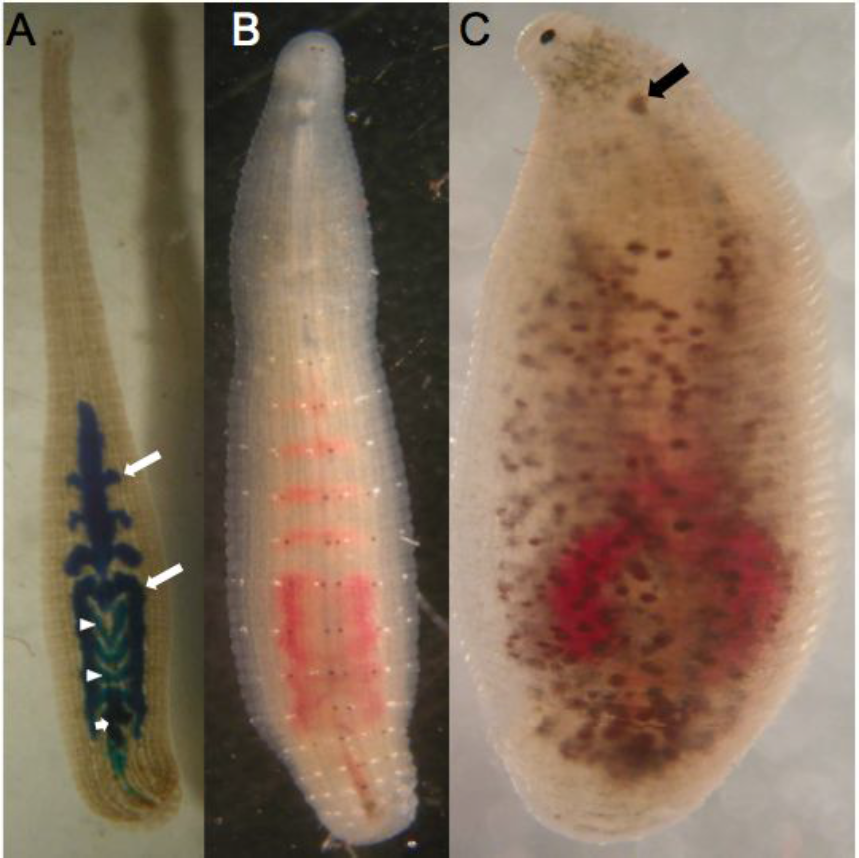
Three species of *Helobdella*. Dorsal views, anterior up, of adult *H. robusta, H. austinensis* and *H.octatestisaca*, respectively, highlighting differences in body wall pigmentation. A) This specimen had fed recently on an artificial food source containing Fast Green dye, which clearly outlines four of the five pairs of large anterior midgut lobes (caecae, long arrows), along with the four pairs of smaller intestinal lobes (arrowheads), and the rectum (short arrow). B) In this animal, which had fed on bloodworms, the crop caecae are labeled red; the central annulus in each segment contains prominent white and brown pigment patches. C) In common with other *H. stagnalis*-like species, this animal bears a chitinous scute (arrow) on the dorsal anterior surface.

**Figure 2.**
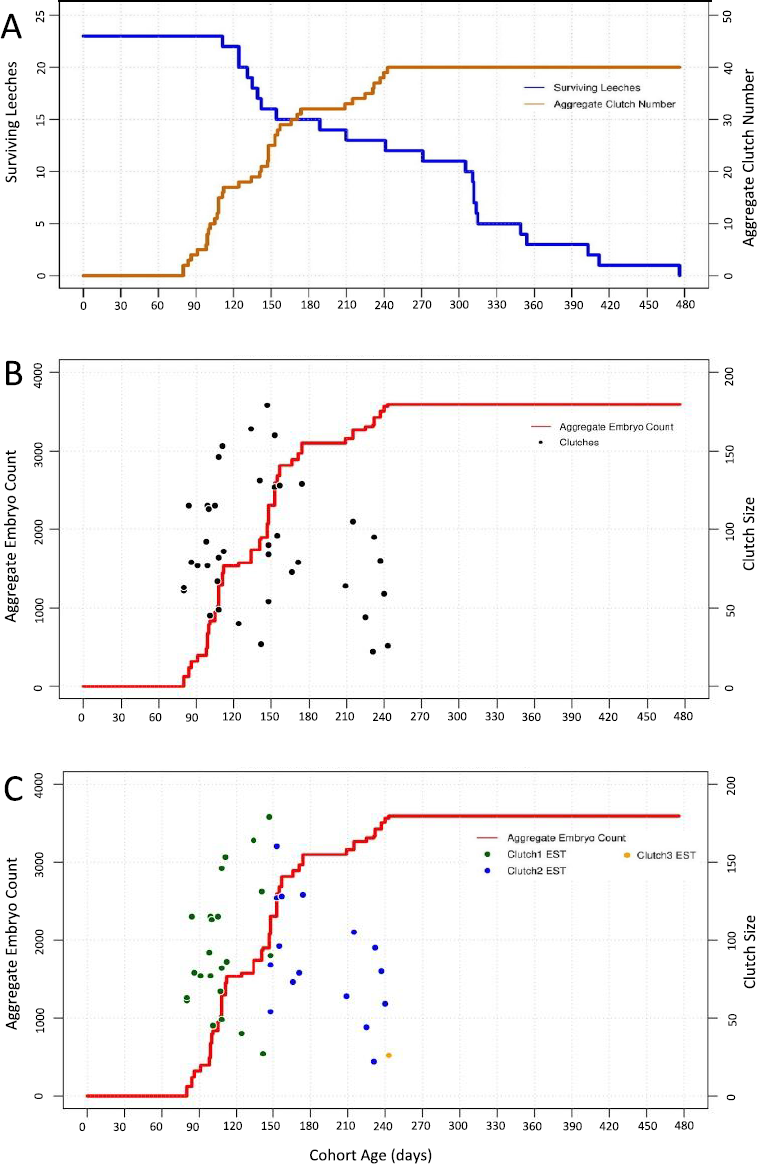
Reproduction in an interbreeding cohort of 23 *H. austensis*. A) Cohort survival (blue, left axis) and aggregate clutch production (orange, right axis) as a function of time, for a cohort of animals fed ad lib on bloodworms. B) Aggregate embryo production (red, left axis); black dots indicate the size (number of embryos, right axis) and deposition date of each individual clutch. C) The same data as in B, except the estimated (EST) assignments of clutches into first, second and third layings are indicated by coloring dots as indicated (see text for details).

**Figure 3.**
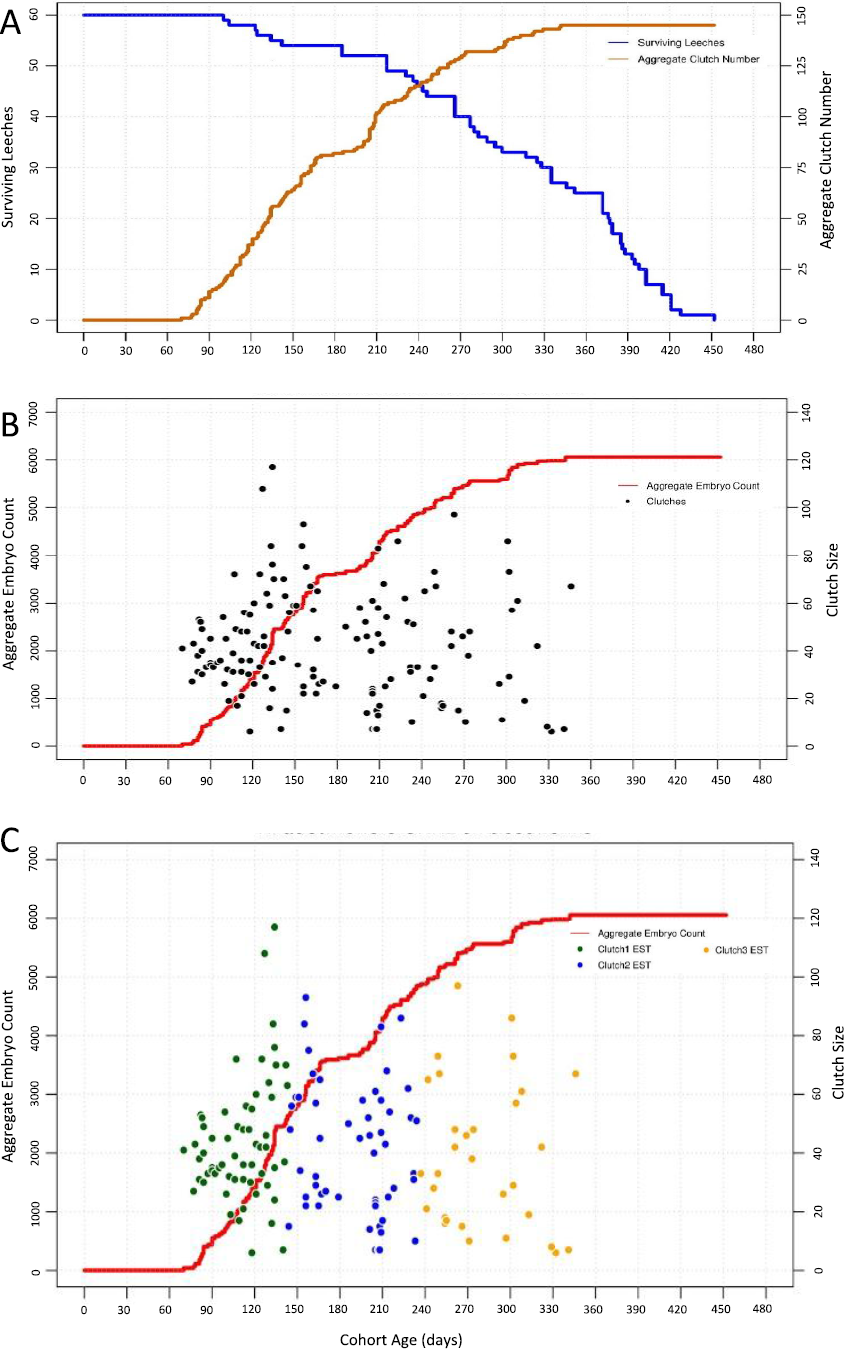
Reproduction in an interbreeding cohort of 60 *H. austensis*. A) Cohort survival (blue, left axis) and aggregate clutch production (orange, right axis) as a function of time, for a cohort of animals fed ad lib on bloodworms. B) Aggregate embryo production (red, left axis); black dots indicate the size (number of embryos, right axis) and deposition date of each individual clutch. C) The same data as in B, except the estimated (EST) assignments of clutches into first, second and third layings are indicated by coloring dots as indicated (see text for details).

The reproductive behavior of individuals within the two cohorts was inferred based on two assumptions: 1) that leeches raised under similar conditions breed in rough synchrony, and 2) that all individuals in the cohort reproduce. Based on these assumptions, we defined the first round of egg laying as beginning with deposition of the first clutch of embryos and ending when the number of clutches deposited equaled the number of animals that had been present when the first clutch was deposited. Similarly, the second and subsequent rounds of reproduction were defined as beginning with the deposition of the next new clutch and ending when the number of additional clutches produced equaled the number of animals that had been present at the beginning of that round of reproduction.

This analysis is subject to various possible errors. For example, if any animals die during the first reproductive round without having produced a clutch of embryos, then what we define as that round would be extended artifactually to include clutches that are actually part of the second round. Conversely, if the temporal spread of reproductive activity is large, the last clutches deposited in the first round of egg laying might be assigned to the second round and vice versa, which would artifactually shorten what we define as the first round. Finally, notwithstanding the fact that all the animals are simultaneous hermaphrodites, we cannot exclude the possibility that some individuals in this experiment were not inseminated and thus failed to deposit zygotes in a given round of reproduction.

Applying this analysis to data obtained for first cohort experiment (starting with 23 individuals) suggests that most animals in the cohort underwent two rounds of reproduction (Figure 2), consisting of 23 and 16 clutches and centered at 109 and 188 days after the birth of the cohort, respectively (Table 2). Based on the assumptions described above, only a single clutch of embryos was assigned to a putative third round of reproduction (Figure 2C, Tables 1, 2). On the other hand, the gap in reproductive activity of the cohort between 175 and 208 days, followed by a cluster of layings between 209 and 240 days, could mean that one of our initial assumptions was in error, and that eight animals underwent a third round of reproduction, from among the 13 surviving at 210 days. In either case, no layings occurred after 243 days, despite the fact that the last individuals in the cohort survived for well over 100 days after the last clutch was deposited. Thus, we concluded that three rounds of reproduction was the maximum observed in this experiment, if the starting assumptions hold true.

The second cohort experiment started with a cohort of 60 *H. austinensis* reared under similar conditions to the first (Figure 3A, B). Interpreting the data from this cohort under the assumptions introduced above again indicates a maximum of three rounds of reproduction, consisting of 60, 54, and 31 clutches and centered at 110, 190 and 273 days after the birth of the cohort, respectively (Figure 3C, Table 2). No further embryos were produced during the last 105 days of the experiment (day 347 through 452), despite the fact that there were 25 surviving individuals at the start of this period. Thus, we again concluded that no individual of *H. austinensis* produced more than three clutches of embryos under these conditions.

### Possible environmental influences on reproductive behavior in *H. austinensis*

Clutch sizes in the two *H. austinensis* cohort experiments varied widely, from 6 to 179 embryos. There was no significant difference between the average size of the inferred first and second clutches *within* either experiment (Table 1). Surprisingly, however, the first two clutches in the first cohort experiment averaged more than twice the size of the corresponding clutches in the second cohort experiment (Table 1). Moreover, the average reproductive capacity in the first experiment (3591 embryos/23 individuals; 156 embryos/individual) was also larger than that in the second experiment (6055 embryos/60 individuals; 101 embryos/individual), despite the fact that more individuals in the second cohort appeared to have laid third clutches of embryos. In any event, two considerations lead us to conclude that these values are conservative estimates of reproductive capacity.

First is the likelihood that some animals in the cohort die without exhausting their reproductive capacity. Premature death could result from disease induced by sub-optimal culture conditions or inadvertent damage while removing embryos for counting.

Another factor is the likely influence of diet on growth and reproduction. *Helobdella* species maintained in our lab are fed frozen chironomid insect larvae (bloodworms), and/or live snails (primarily *Lymnaea* and *Physa*); the three species of *Helobdella* studied here exhibit different dietary preferences. *H. octatestisaca* prefers bloodworms; snails placed in their bowl survive for many days as long as bloodworms are provided. In contrast, *H. robusta* exhibits a strong preference for snails; we have not succeeded in maintaining this species on a pure bloodworm diet. Finally, *H. austinensis* feed and breed readily on either bloodworms or snails. While a systematic investigation of the links between diet and reproductive capacity was beyond the scope of the present work, this species grows much larger when fed snails; *H. austinensis* fed with excess bloodworms seldom exceed 40 mg in size (Rachel Kim, personal communication), and the maximum clutch size for animals on a bloodworm diet was 179 embryos (Table 1); in contrast, snail-fed individuals can grow to more than 120 mg and produce single clutches of over 200 embryos (Shinja Yoo, personal communication).

### Breeding cohorts of *H. robusta* exhibit clustered bouts of reproduction

The indirect conclusion that *H. austinensis* exhibits a maximum of three bouts of reproduction was similar to our observations based on direct observation of reproductive behaviors of self-fertilizing individual *H. octatestisaca* individuals, but markedly different than for what we observed for self-fertilizing individual *H. triserialis* (up to five layings) and *H. robusta* (up to eight egg layings) (Supplemental Table 1, Tables 1 and 2). The strength of these inter-species comparisons is limited, however, by the differences in the experimental conditions--some of the observed differences might reflect differences between animals in interbreeding cohorts versus self-fertilizing animals in isolation.

It is obviously not possible to observe the reproductive behavior of individual *H. austinensis* in isolation. Thus, to compare the reproductive behavior of a cohort of leeches to that of isolated conspecifics, we re-collected *H. robusta* from the type location (Kutschera et al. 2013), confirmed the species identity by CO1 sequencing (see Materials and Methods), and then carried out a cohort breeding study starting with animals originating as a single clutch of embryos. The experiment was carried out as described above for *H. austinensis* except that the *H. robusta* were fed on their preferred diet of live, lab-reared snails, as for the experiments on isolated, self-fertilizing *H. robusta.*

Starting with a cohort of 48 animals, a total of 7304 embryos from 149 clutches were produced ranging in size from 11 to 114 embryos (Tables 1 and 2; Figure 4A, B). Casual inspection of the data suggested several differences between reproductive behavior in *H. austinensis* and *H. robusta*: 1) bouts of reproductive activity in the *H. robusta* cohort were more tightly clustered than for *H. austinensis* in both timing and clutch size; 2) there appeared to be five such bouts, and: 3) there was an apparent correlation between clutch size and reproductive episode, with the third and fourth clutches being the largest. There was no significant difference in the egg-to-egg generation time between the two species under these conditions (Table 2).

**Figure 4.**
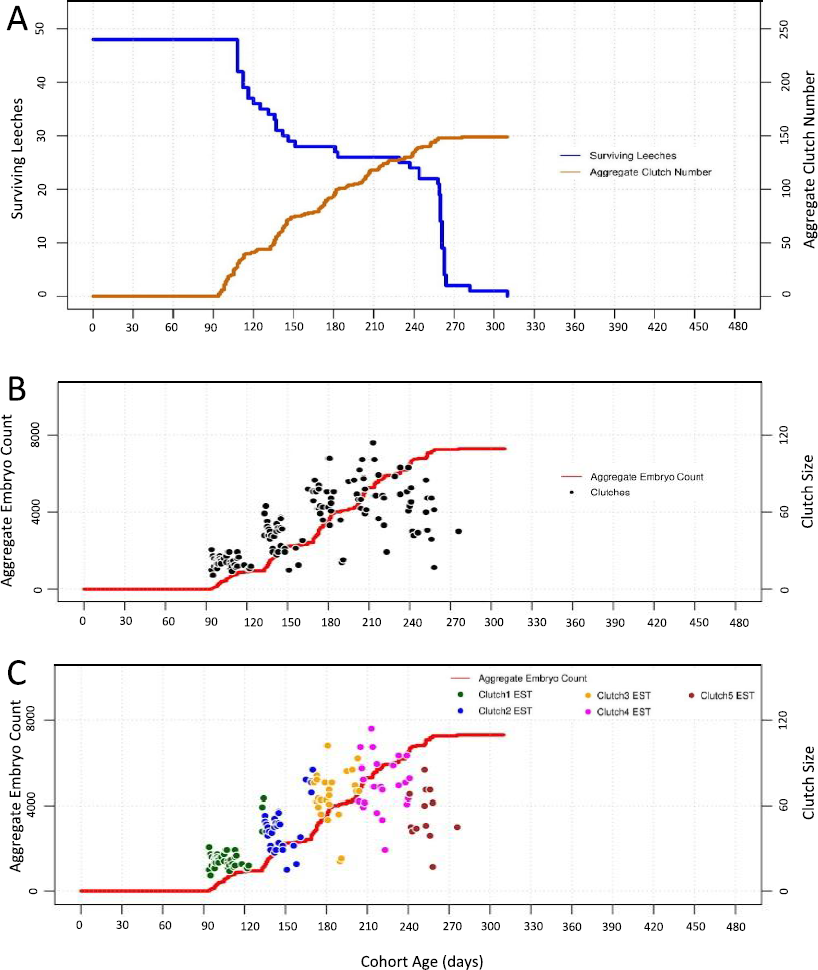
An interbreeding cohort of 48 *H. robusta* exhibits clustered bouts of reproduction. A) Cohort survival (blue, left axis) and aggregate clutch production (orange, right axis) as a function of time, for a cohort of animals fed ad lib on snails. B) Aggregate embryo production (red, left axis); black dots indicate the size (number of embryos, right axis) and deposition date of each individual clutch. C) The same data as in B, except the estimated (EST) assignments of clutches into first, second and third layings are indicated by coloring dots as indicated (see text for details).

Analyzing the reproductive behavior data under the same assumptions as for *H. austinensis* suggested that the cohort reproduced in five clusters of 48, 34, 28, 26 and 13 layings, respectively (Figure 4C; Tables 1 and 2). There was a sharp drop in population after the fifth cluster of reproductive activity, and the last surviving leech in the cohort died only 34 days after the last clutch of embryos was laid. This sharp decline contrasts with the gradual decline and extended post-reproductive survival of individuals in the two cohort experiments for *H. austinensis* (cf Figures 1 and 2). In light of these observations, one interpretation is that the *H. robusta* cohort died off prematurely, either due to parasites picked up from the snails, or to other, unknown factors. Problems of colony decline and extinction have been noted by ourselves and others for *H. triserialis* and *H. robusta* (M. Shankland, personal communication; D-H Kuo, personal communication), and are responsible for the shift to using *H. austinensis* as a more lab-tractable species for study.

### Reproduction parameters derived from isolated *H. robusta* do not predict the clustered bouts of reproductive activity seen in breeding cohorts

To compare the reproductive behaviors of *H. robusta* in isolation and in cohorts, we first plotted the combined reproductive data from 16 isolated individuals, comprising a total of 75 clutches, to ask how well the resultant “pseudo-cohort” data recapitulated the reproductive behavior of the actual cohort (Figure 5A, B). For this dataset, we could also determine how well the reproductive behaviors inferred using the methods applied to the *H. austinensis* cohorts (Figure 5C) match the actual behavior of the individuals comprising the pseudo-cohort (Figure 5D).

**Figure 5.**
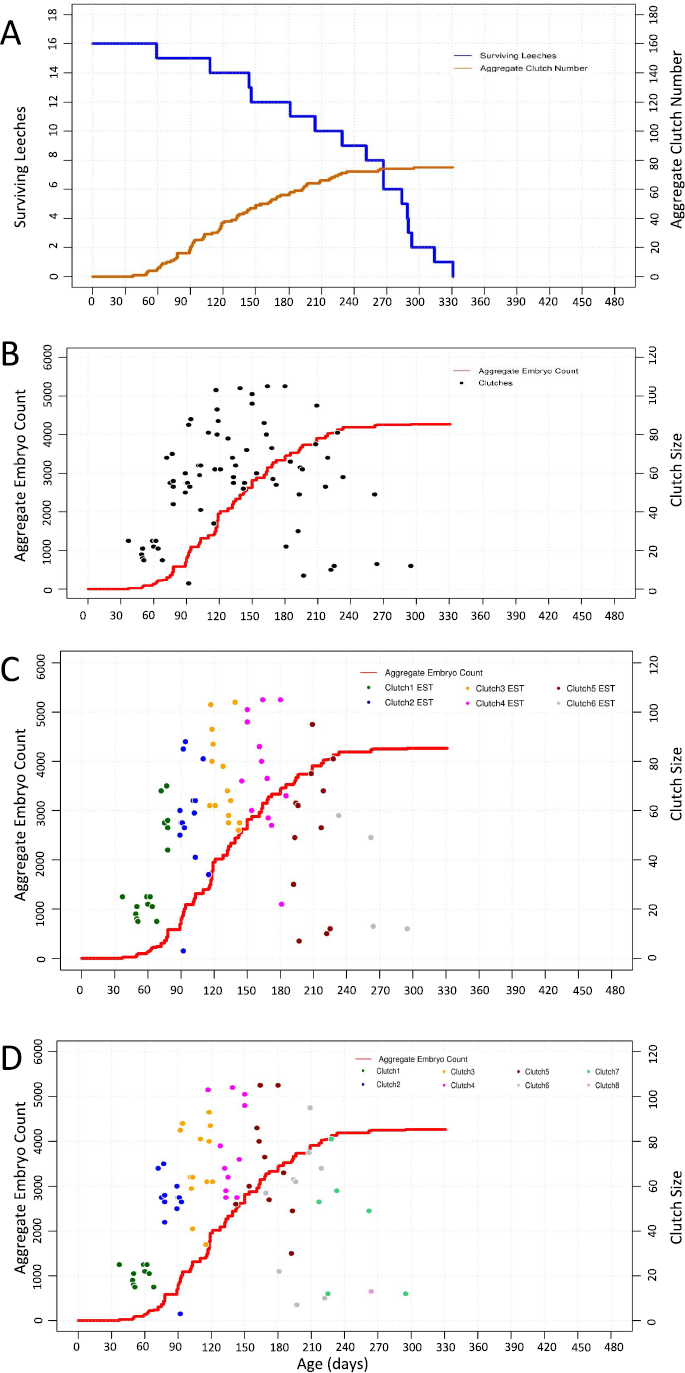
Clustered reproductive activity by *H. robusta* cohort is not predicted from reproductive parameters derived from isolated individuals. A-C) A pseudo-cohort was created by graphing the aggregate data from 16 animals reared in isolation as for the true cohorts in Figures 2-4. D) For comparison, the actual clutch groupings are denoted using the same color scheme. Note that some clutches inferred as being part of the second round of reproduction actually occurred in the first round, and that only six rounds of reproduction were inferred, whereas the true value was eight.

Surprisingly, the pseudo-cohort data (Figure 5) differ from the cohort data (Figure 4) in at least two ways. First, the onset of reproductive activity in isolated animals was markedly earlier than in the cohort experiment. Among animals raised in isolation, the earliest reproductive episode occurred just 37 days into the life of the parent and all 16 individuals had completed their first round of reproduction by 68 days (Table 2); indeed, most animals reared in isolation had completed their *second* round of reproduction before the first cluster of reproductive activity among the cohort animals. Second, the discrete clusters of reproductive activity in the cohort were largely absent from the pseudo-cohort, especially after the first bout of reproduction. A third difference is that the cohort seemed to undergo a maximum of five rounds of reproduction, whereas isolated individuals deposited up to eight clutches of embryos. As mentioned above, however, we cannot exclude the possibility that the cohort population died off before exhausting its reproductive capacity. Moreover, comparing the inferred clutch groupings (Figure 5C) with the actual clutch groupings (Figure 5D) shows that the inference method predicted a wider temporal distribution of clutch deposition times and a reduced number of egg layings (six) relative to the actual data (eight).

### Computer simulations of reproductive activity

As a further inquiry into the apparent difference in reproductive activity between isolated individuals and an interbreeding cohort of *H. robusta*, we modeled cohort breeding data through development of a Monte Carlo simulator with an automated graph plotter (details of the program and instructions for use are available upon request). In one set of simulations, the program probabilistically generated values for time-to-first-clutch, clutch size, inter-clutch intervals and survival times, using life history data derived from the pool of 16 *H. robusta* reared in isolation. In a second set of simulations, the corresponding values were generated using parameters inferred from the cohort of 48 *H. robusta*. The simulations also allowed us to match the size of the simulated cohorts to those of experimental cohorts.

The first set of simulations, based on parameters derived from individuals raised in isolation, accurately reproduced behavioral activity of the pseudo-cohort as expected (compare Figures 5D and 6A), but failed to fully reproduce that of the true cohort (compare Figures 4C and 6B). The first bout of reproductive activity was clustered (indicating low variance in the zygote-to-first-clutch generation time), but occurred much earlier than we observed in the actual cohort experiment. Repeated simulations for cohorts of either 16 or 48 individuals failed to produce the tightly clustered bouts of reproduction that had been observed throughout the actual cohort experiment, indicating that the variance of the inter-clutch intervals was higher for the individuals than for the cohort. Also as expected, the overall productivity of the simulated cohort of 48 animals was higher than was observed for the actual cohort of 48 animals.

The second set of simulations used parameters inferred from the cohort of 48 *H. robusta*. As expected, these simulations performed better in reproducing the zygote-to-first clutch generation time for the cohort, but still failed to capture the temporally clustered bout of reproduction observed in the true cohort (compare Figures 6D and 4C), indicating that the variance of the inferred inter-clutch intervals was higher than that of the actual inter-clutch intervals. We interpret this discrepancy as revealing that one or more animals counted as present at the beginning of a bout of reproduction either died or otherwise failed to reproduce, so that in counting up the clutches and assigning them to what we defined as one round of reproduction actually included clutches that were part of the subsequent round. As expected, simulating cohorts of 16 animals using parameters inferred from the cohort of 48 animals predicted lower productivity and also a tighter temporal clustering of reproductive activity than was obtained with the pseudo-cohort of 16 animals (compare Figures 6C and 5D).

**Figure 6.**
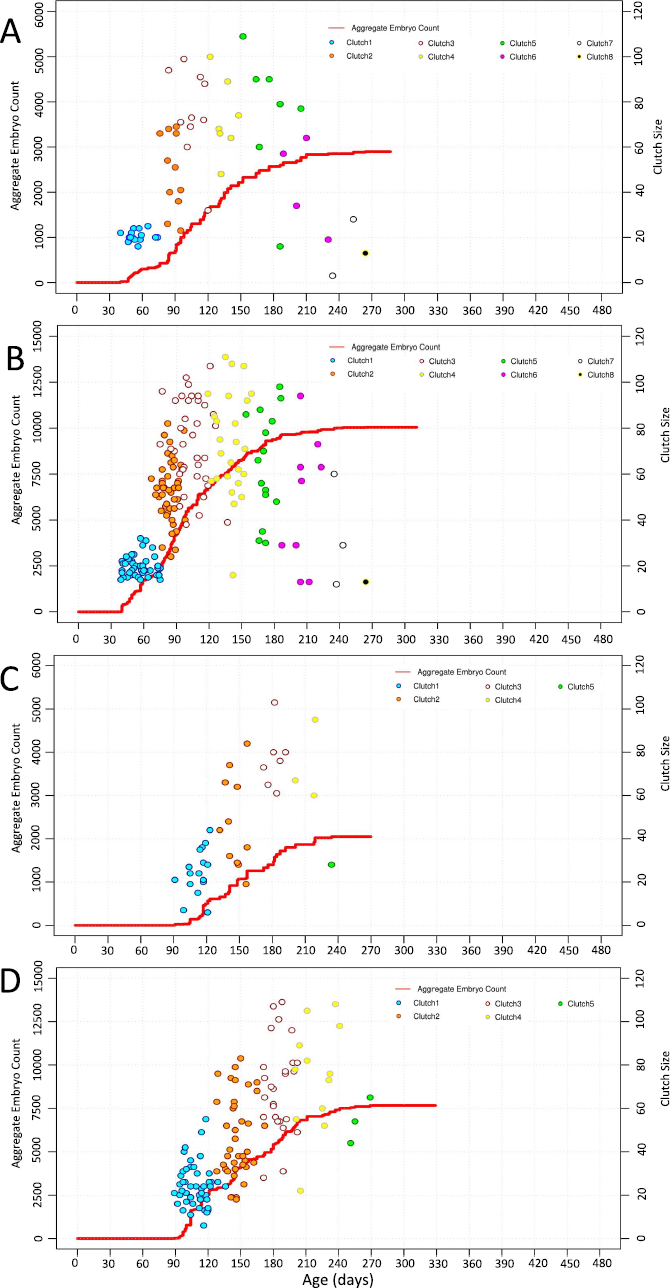
Monte Carlo simulations of reproductive activity in *H. robusta.* A, B) Simulated reproductivity of 16 and 48 animals, respectively, using parameters from 16 self-fertilizing animals raised in isolation. C, D) Simulation of 16 and 48 animals, respectively, using parameters inferred from the cohort of 48 interbreeding animals. Note that none of these simulations capture the temporal clustering observed in the experimental cohort of 48 animals.

Thus, we conclude that the parameters for reproductive behavior of isolated *H. robusta* cannot account for the coordinated reproductive behavior exhibited by cohorts of interbreeding individuals.

## DISCUSSION

The work presented here examines the reproductive life histories, under laboratory conditions, of three glossiphoniid leech species in the genus *Helobdella*: *H. austinensis, H. octatestisaca* and *H. robusta.* This work complements a previous description of reproductive behavior for a fourth species, *H. triserialis* (Wedeen et al, 1990).

One major distinction in reproductive life strategies for both plants and animals is whether individuals of a species reproduce only once (semelparity) or more than once (iteroparity) before dying. Semelparity has been documented for several glossiphoniid leech species including *Alboglossiphonia polypompholyx* (El-Shimy and Davies 1991), *Marsupiobdella africana* (Van der Lande and Tinsley 1976), *Theromyzon cooperei* (Oosthuizen and Fourie 1985), *T. rude* and *T. tessulatum* (Wilkialis and Davies 1980). In contrast, iteroparity holds for several other leech species, e.g., the medicinal leech (*Hirudo medicinalis*; Davies and McLoughlin 1996).

All sexually reproducing clitellate annelids (oligochaetes and leeches) are simultaneous hermaphrodites, but self-fertilization is rare. Self-fertilization has been reported for the piscolid species *Zeylanicobdella arugamensis* (Kua et al. 2010) and for the glossiphoniids *Clepsine marginata* (Whitman, 1878), *Helobdella triserialis* (Wedeen et al. 1990), and *H. papillornata* (Tan et al., 2004)--this species is apparently identical to the previously described *H. europea* (Pfeiffer et al. 2004). We have previously speculated (Cho et al. 2014) that a capacity for self-fertilization may contribute to the species richness of the genus *Helobdella* (see below)--if self-fertilization can rescue genomic rearrangements that would result in otherwise infertile individuals, it would result in reproductive isolation of nascent species without significant changes in habit or habitat (sympatric speciation).

### Interspecies differences in reproductive life history

Based on the work presented here, iteroparity and the capacity for self-fertilization appear to be the rule in the genus *Helobdella*, but our work reveals a number of species-specific differences in reproductive capacity. Under similar laboratory conditions, three of the species we have studied, *H. octatestisaca*, and *H. robusta* in the present work, and *H. triserialis* in previous work are all self-fertile. Studied as isolated individuals, there were clear differences in reproductive behavior among these three species: *H. octatestisaca* individuals never produce more than three clutches of cocoons, *H. triserialis* routinely produce five clutches, but never more, and *H. robusta* produce up to eight clutches (Tables 1-2; Supplemental Tables 1-3).

In previous studies, reproductive life histories for *Helobdella* have been inferred from systematic measurements of size and reproductive status of wild-caught animals at different times of year for species identified as *H. stagnalis* in Wales (Learner and Potter, 1974; Murphy and Lerner, 1982;) and in Tunisia (Romdhane et al. 2017). The conclusion of these studies was that *H. stagnalis* undergoes two rounds of reproduction in the field.

*Helobdella* appears to be a speciose genus compared with other groups of glossiphoniid leeches (Oceguera-Figueroa et al., 2010). As has proven to be the case for some other widespread taxa, molecular sequence analyses have led to identification of cryptic species (Bely 2006). Thus, until recently, all leeches bearing a nuchal scute (Figure 1C), were classified as *H. stagnalis*, but there now appear to more than a dozen such species (Saglam et al. 2018; Beresic-Perrins et al. 2017; Oceguera-Figueroa et al. 2010). These *H. stagnalis*-like species do not form a distinct clade based on cytochrome oxidase 1 (CO1) sequence alone (Oceguera-Figueroa et al. 2010), but this may reflect the limited number of informative sites in the CO1 sequence. Based on its CO1 sequence, the *H. stagnalis*-like species we studied here is *H. octatestisaca*, originally described from Taiwan but apparently introduced there from Mexico (Lai et al. 2009); its presence in California could represent a broader natural range than previously thought, perhaps thanks to migratory waterfowl or introduction by humans. Whether the difference between producing two clutches (for the morphologically defined *H. stagnalis* in Wales and Tunisia) or three clutches (for *H. stagnalis*-like species we identify as *H. octatestisaca*) reflect genuine inter-species differences or differences between laboratory and field conditions remains open.

### Inferring reproductive parameters from interbreeding cohorts

Given that four species of *Helobdella*, representing different branches of the clade are self-fertile, it seems parsimonious to assume that this capacity is ancestral within the species. Thus, we were surprised to find that *H. austinensis* appears incapable of reproducing by self-fertilization.

To infer the reproductive behavior of individuals in this species, we followed two interbreeding cohorts throughout their entire lifespans, and concluded that this species also produces a maximum of three clutches of embryos (Figures 2 and 3). These inferences were drawn based on the basic assumption that the reproductive behavior of animals within the cohort is approximately the same. We note that inter-animal variations in the distribution of surface markings such as papilla and pigment cells in *H. austinensis* should make it possible to identify and distinguish individual animals (Kutschera et al., 2013). In principle, such morphological heterogeneity could make it possible to track the reproductive behavior of individuals within a cohort directly. Such an undertaking was beyond the scope of the present work, however.

Comparing the reproductive parameters of the two cohorts of *H. austinensis* revealed that the temporal features of reproductive activity were well conserved between the two experiments, as judged by both the distribution of egg-to-egg generation times and the inferred inter-clutch intervals (Table 2). In contrast, the reproductive capacity differed markedly between the two cohorts, averaging 156 zygotes per individual in the smaller cohort (starting with 23 individuals), compared with only 101 zygotes per individual in the larger cohort (starting with 60 individuals). This difference cannot be explained by differences in cohort survival--in both experiments, many animals survived for weeks after the cessation of reproductive activity, and the larger cohort deposited more cocoons overall; rather, the first and second clutches for the smaller cohort averaged more than twice the size of those in the larger cohort.

Both cohorts were raised in the same size of container. Thus, the population density was higher for the larger cohort than for the smaller one. In this context, the difference in the progeny produced is consistent with theoretical predictions and experimental observations on the population density-dependence of sperm competition and reproductive resource allocation in simultaneous hermaphrodites (Schärer 2009; Schärer and Pen, 2013; Cannarsa and Meconcelli, 2017). In brief, and taking the extreme case of a single self-fertile hermaphrodite, the optimal reproductive strategy for such an individual would be to make as many eggs as energetically feasible, and restrict sperm production to the bare minimal required to fertilize those eggs. In contrast, as the population density increases, and thus the probability of interbreeding instead of self-fertilization, it is advantageous to make more sperm, in the expectation of being able to fertilize eggs from another individual, and fewer of the energetically more costly eggs, which are more likely to be fertilized by another individual.

These ideas have been tested in various species, including experiments with the leech species *H. papillornata*/*H. europea* (Tan et al., 2004). Using total volume of testisacs and eggs as proxies for investment in sperm and eggs, respectively, these authors found that normalized testisac volume increased with increasing group size, but that egg volume did not. Our experiments do not permit statistical tests, but do suggest that for iteroparous species, measuring differences in overall egg production during the reproductive life of the individual, rather than at a single time point, might also reveal plasticity in maternal as well as paternal investment at different population densities.

### Intra-species differences between reproduction by self-fertilization and interbreeding

To look for differences between the reproductive behavior of self-fertilizing and interbreeding individuals within the same species, we also followed the reproductive behavior of a *H. robusta* cohort. This experiment yielded three noteworthy results.

First, animals in the cohort exhibited a significant delay in the onset of reproduction compared to individuals reared in isolation (107.4 +/- 10.7 days vs. 56.3 +/- 8.7 days). This result differs from observations on *H. papillornata* by Tan et al. (2004), who reported that “Self-fertilization is possible, because isolated individuals have produced offspring in the laboratory, but our observations suggest that individuals resort to self-fertilization only after a long period in which no partners could be found.” Notwithstanding these observations of delayed reproduction in isolated *H paillornata*, it would also seem reasonable for isolated self-fertile animals to initiate reproductive activity as soon as possible, to increase the population size--thereby increasing the probability of surviving the population bottleneck, and also enabling dispersal to increase the chances for encountering other conspecifics for subsequent interbreeding.

Second, the reproductive capacity of *H. robusta* in the cohort (averaging 149 zygotes per individual) was much lower than those raised in isolation (averaging 267 zygotes per individual). In contrast to our observations for *H. austinensis*, however, the difference in between self-fertilizing and interbreeding *H. robusta* arises from differences in the number of clutches produced, and not in clutch size. Self-fertilizing animals produced an average of 3.7 clutches per individual, with an observed maximum of eight, whereas the cohort-reared animals produced an average of 2.4 clutches each, with an inferred maximum of five. The difference in reproductive capacity is again consistent with the predictions of reproductive resource allocation theory. In this case however, as noted above, it is also possible that more animals in the cohort died before exhausting their reproductive capacity.

A final intriguing difference between the reproductive behavior of *H. robusta* in isolation, as opposed to an interbreeding cohort, is the clustering of reproductive episodes among individuals in the cohort. Monte Carlo simulations confirm that this clustering reflects a tightly distributed timing of reproductive episodes which cannot be explained based on the reproductive behavior of animals in isolation (Figs. 4-6). Precisely synchronized reproduction is well-known in certain marine polychaetes, providing the advantages of increased probability for encountering mates and overwhelming predators by mass producing spawn (Fischer 1999; Pamungkas and Glasby 2015), but has not been noted for leeches.

A possible model to explain this clustered reproductive behavior starts with the notion that maternity is much more costly than paternity for glossiphoniid leeches, whose reproduction involves a large maternal investment: first, cross-fertilization is by traumatic insemination, in which spermatophores implanted into the body wall of the partner digest their way through the multiple layers of the body wall before releasing sperm into the coelom (Sawyer 1986); in addition, glossiphoniid leeches make large, yolk-rich eggs, brood their embryos in cocoons attached to the ventral aspect of the parent and carry the juveniles with them to the first one or more feedings. Given the high cost of the maternal role for these hermaphrodites, it would seem advantageous for individuals in a cohort to retard maturation of their eggs until others in the cohort are susceptible to being sperm acceptors as well as sperm donors, thereby balancing out the physiological costs of maternity with the advantages of paternity.

We speculate that this model could account for both the delayed onset of reproductive activity in the cohort relative to the isolated individuals, and for the clustered reproductive activity exhibited by cohorts of *H. robusta* relative to individuals. In any case, we conclude that *Helobdella* species provide a phenomenologically rich, experimentally tractable resource for further investigations of reproductive life history strategies.

